# Pesticide-exposed bees fail to thermoregulate leading to cold colonies with consequences for offspring development

**DOI:** 10.64898/2025.12.03.692033

**Authors:** Kieran E. Storer, Susan E.G. Hawthorne, Connor Lovell, Ferozah Mahmood, Daniel Kenna, Tianqi Zhu, Peter Graystock, Richard J. Gill

**Affiliations:** Georgina Mace Centre for the Living Planet, Department of Life Sciences, Imperial College London, Silwood Park campus, Ascot, SL5 7PY, UK; University of Oxford, Department of Biology, South Parks Road, Oxford, OX1 3RB, UK; Kings College London, Institute of Zoology Strand, London, WC2R 2LS, UK

**Author notes:** joint first authors (Kieran E. Storer, Susan E.G. Hawthorne).

## Abstract

For many organisms, effective thermoregulation is needed to cope with changing environmental temperatures. But if exposure to pesticides in the environment were to impair this homeostatic process, reproduction and population viability could be at risk. Focusing on an important insect pollinator, we conducted three complementary experiments exposing bumblebees to a pesticide under different temperature challenges and measuring the impacts on thermoregulatory ability and brood development. First, we reveal that pesticide-exposed individuals cannot maintain a stable thorax temperature, especially at lower temperatures. Second, reductions in body temperature are accompanied by behavioural changes and that pesticide-exposed colonies fail to maintain appropriate brood temperatures. Third, such collective impairment on brood thermoregulation (not the pesticide toxicity to offspring *per se*) leads to delayed pupal development and reduced adult population growth. Our study provides a valid mechanistic explanation for why terrestrial insects requiring brood thermoregulation have declined. With frequent extreme weather events forecasted, our findings have concerning implications for how populations will adequately persist and grow under current pesticide-use regimes with ramifications on pollination services.

## 1. INTRODUCTION

Environmental pesticide exposure poses potential health risks to a range of insects that perform vital ecosystem functions (1–3). Whilst mortality from acute exposure is rare (4), a culmination of sublethal effects on physiology and behaviour may translate to population-level impacts (5–7). For the ecologically and economically important social insects (i.e., ants, termites, species of bees and wasps), theory predicts that subtle impairments to individual function could build up to result in colony collapse (5, 8). Direct empirical support for this, however, remains surprisingly limited, as pesticide exposure studies do not commonly test across multiple levels of biological organisation (9–11). For instance, whilst pesticide exposure has been associated with lower colony productivity (12–17), it is unclear how this has emerged from pesticide induced changes to individual behaviour and physiology. Identifying this link is important for revealing causative pathways and determining how to engineer and target biological solutions (18). Furthermore, when quantifying sublethal effects, we must consider the environmental context dependency of pesticide toxicity (19). For instance, quantifying how pesticide impacts are modulated by different ambient temperatures is crucial to forecast pesticide risks across climatic regions and under climate change (20–23).

Adaptive strategies to buffer climatic challenges require adult individuals to regulate their own metabolic activity for attaining appropriate body temperature and thermoregulate the nest/brood for successfully rearing offspring (24–26). This is important for social insects with permanent nest sites, as they must deal with unfavourable environmental temperatures that frequently arise. Given that certain pesticides, such as neonicotinoids, have been reported to affect molecular and cellular processes involved in energy metabolism and mitochondrial function (27–31) and impair non-flight thermogenesis (32, 33), exposure to such chemicals might impact thermoregulatory behaviours and jeopardise colony adaptability to changing environmental temperatures (10). If such impacts on individual behaviour(s) translate to decreased colony function and productivity, then populations may be at risk from the combined threats of increased pesticide usage (34) under climate change (35). Thus, with bee colonies increasingly experiencing extreme weather events and being active later in the year (36), quantifying temperature dependent pesticide impacts on bee thermal responses represents a research priority.

Here we investigate a mechanism by which exposure to a systemic and cholinergic pesticide leads to temperature dependent impairments of bee colonies. We focused on a neonicotinoid insecticide (imidacloprid) for two key reasons: i) despite recent restrictions, neonicotinoids remain one of the most used insecticide classes globally (37, 38) with bee exposure via contaminated nectar and pollen seeming to be a common occurrence along with residue accumulation inside colonies (9, 39, 40); ii) by building from previous work (10, 21) we can develop a comprehensive understanding and set of first principles for how pesticides that share similar modes of action can impair organisms across different levels of biological organisation (e.g. genes to ecosystem; that can improve understanding of both past responses and future risks (41)).

Conducting three complementary experiments involving sublethal pesticide exposure assays under different controlled temperatures, we tested the hypothesis that higher demands under cooler temperatures on individual thermogenesis leaves pesticide-exposed colonies less able to maintain nest temperature with impacts on brood development (Fig. 1). For experiment 1 we tested whether exposed individual bumblebee workers can maintain body temperature (Fig 1. arrow a; Fig. S1). For experiment 2, we linked how exposure altered the collective behaviours of individuals, with a focus on brood attendance, and tested how this was associated with brood temperature and colony adult population growth (Fig 1 arrows b & c; Fig. S1). For experiment 3, we tracked (*in vitro*) development of isolated early-stage pupae removed from control and exposed parental colonies at a low and high temperature regime that reflected end of assay brood temperatures measured in experiment 2 (Fig 1 arrows c & d).

**Fig. 1.**
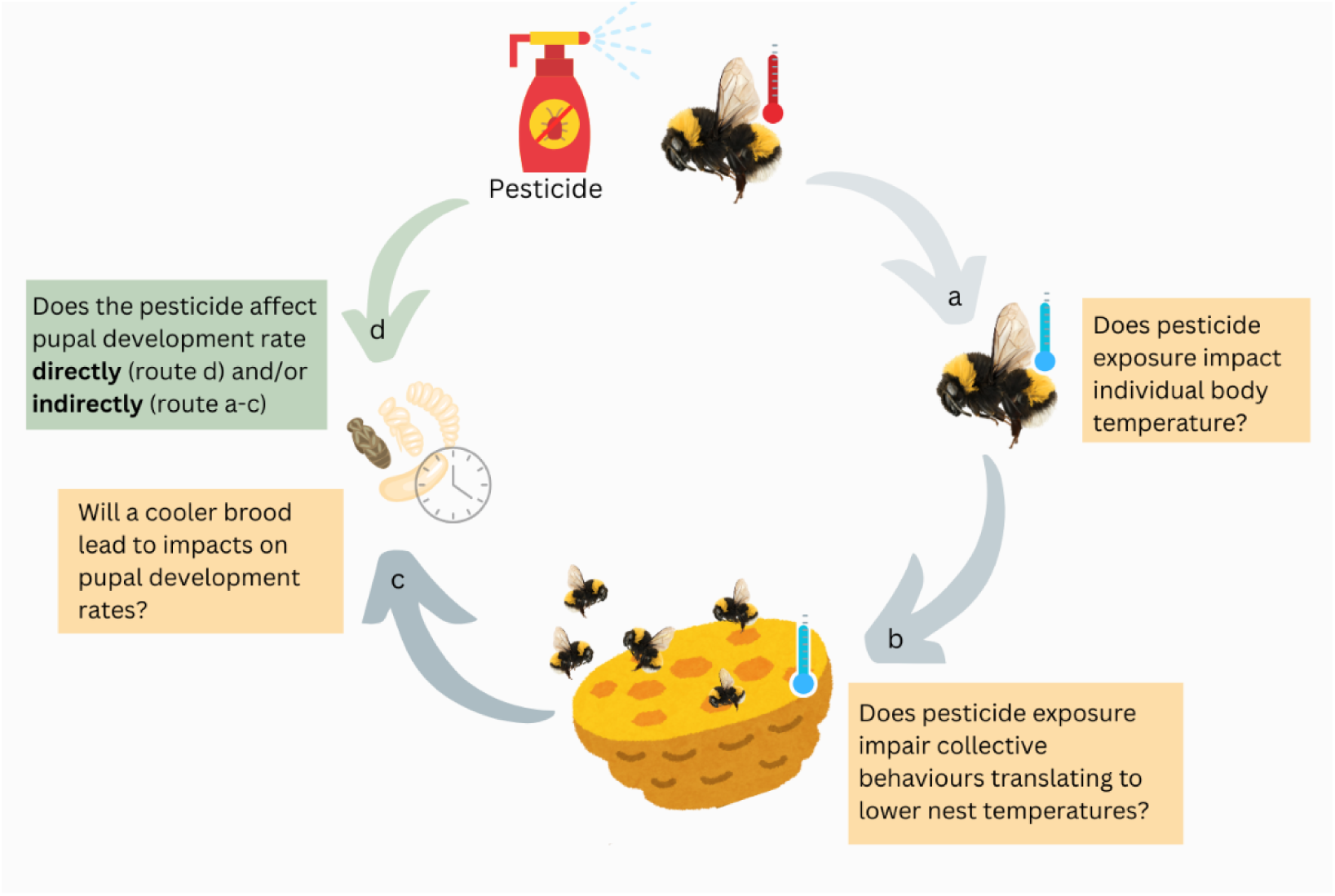
Conceptual diagram outlining our multi experimental approach and hypotheses to be tested concerning how pesticide exposure affects bumblebee individual and colony thermoregulation with impacts on pupal development. We test how pesticide exposure affects individual body temperature (arrow a, experiment 1), how this translates to colony behaviours and nest thermal homeostasis (arrow b, experiment 2), and if this indirectly culminates in impacting pupal development rates (arrow c, experiment 2 & 3). We further distinguish whether the impact on pupae is from the indirect route of impaired thermoregulation (yellow boxes and grey arrows a-c) or if direct pesticide toxicity can also explain our results (green box and arrow d, experiment 3). We test each hypothesis across a range of environmental temperatures.

## 2. RESULTS

### 2.1 Experiment 1: Temperature dependent effects of pesticide exposure on individual thorax temperature profiles

Control bees held a stable relative thermal profile at all three ambient temperatures over the trial, maintaining a mean thorax surface temperature of 28.2°C when at 15°C ambient (var=6.6), 30.6°C at 20°C ambient (var=4.0), and 32.3°C at 25°C ambient (var=1.8). Pesticide-exposed bees, however, showed declines in thorax temperature and higher variation between individuals. For general comparison, with the effect of time not considered, pesticide-exposed bees had a lower mean thorax temperature of 26.1°C at 15°C ambient (var= 27.3; W = 14342, p <0.01) and 29.8°C at 20°C ambient (var= 8.0; W = 13413, p <0.01). In contrast, when at 25°C ambient pesticide bees on average had a higher thorax temperature, and again with higher relative variation (mean: 32.7°C, var 2.281; W = 2823.5, p = 0.026).

Pesticide effects were, however, time dependent with initially similar relative thorax temperature to control bees, before declining significantly as the trial progressed (*pesticide x time*: t = -4.62, p < .001; Fig. 2; Table S8). This effect was greater at the lowest (15 °C) ambient temperature (*pesticide* x *temperature*: t = 2.44, p = 0.015). Intriguingly, at 25°C, the thorax temperature of pesticide-exposed bees was on average higher compared to control bees in the initial 2-3 hours of the experiment, before converging with control bees towards the end of the trial. Bees that consumed less nectar substitute relative to their body size could not maintain their thorax temperature as high (t = -3.34, p < .001). Furthermore, pesticide-exposed bees tended to consume more food relative to their body size compared to unexposed bees when at 15°C (W = 15179, p-value < 0.001), presumably attempting to raise their body temperature. Indeed, when at 20°C consumption did not differ between pesticide and control (W = 12786, p-value = 0.067), and became less in the pesticide group at 25°C (W = 1741, p = 0.007).

**Figure 2.**
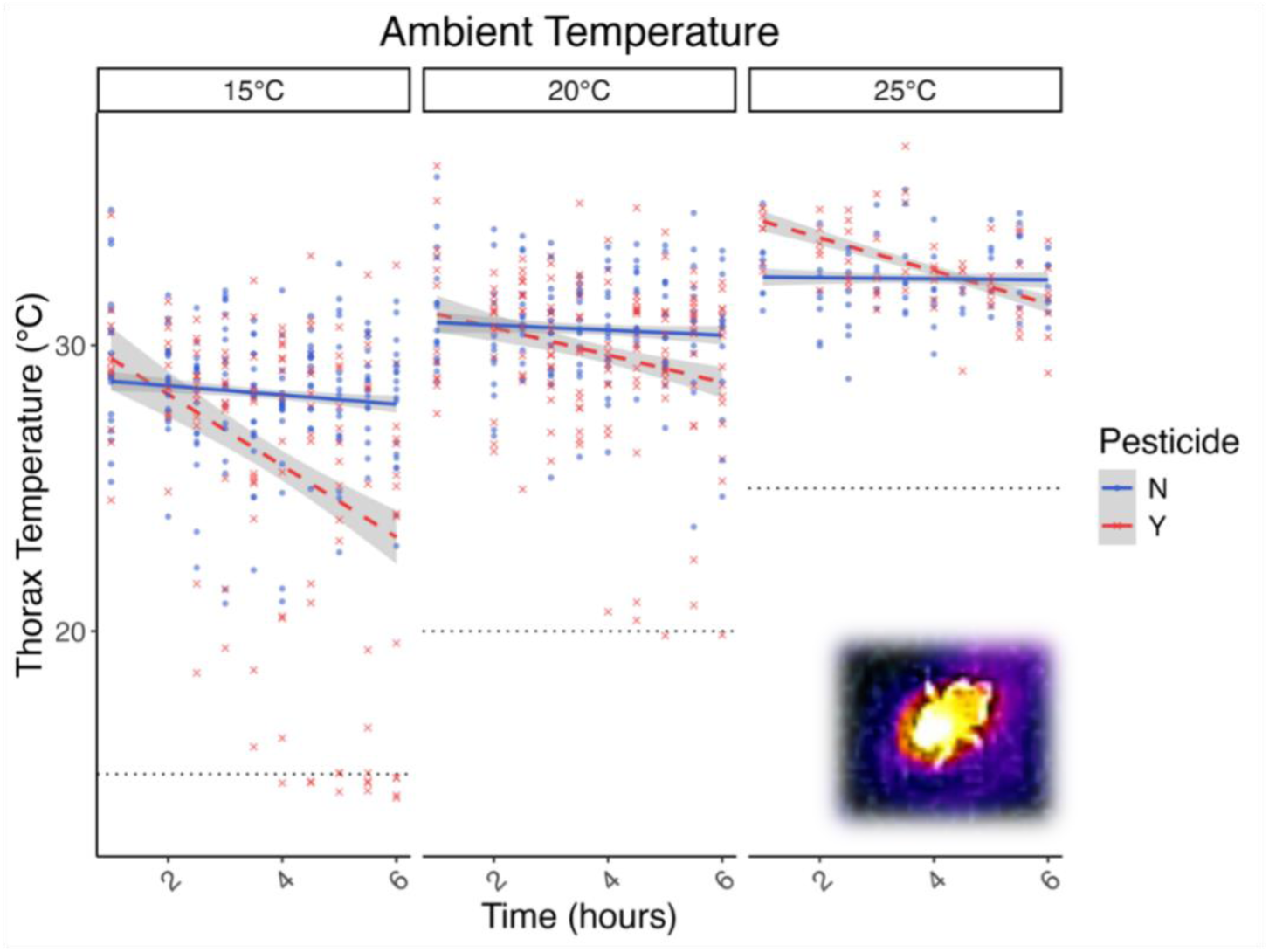
Pesticide-exposed bumblebees are unable to maintain a consistent thorax temperature (experiment 1). Bee thorax surface temperature for pesticide-exposed (red crosses and dashed line) and control (blue circles and solid line) bees over the 360-min trial at the three ambient temperatures (15, 20 and 25°C, also shown as horizontal black dotted reference lines). Data lines represent estimates from the GLMER model with grey ribbons showing 95% CIs, and raw datapoints shown in the background (n = 127 individual bees across treatments). Inset is a thermal image of an individual bumblebee.

### 2.2 Experiment 2: Temperature dependent pesticide effects on worker activity, brood care behaviour and cohort brood temperature

#### 2.2.1 Worker activity

On day-1 of the trial, over half of the bees across experimental groups were classified as active and in physical contact with the brood (Table S9). By day 5, however, the behaviour between pesticide and control groups developed to be distinctively different, with 93.3% and 72.0% showing a behaviour classified as inactive in the pesticide group when at 19°C and 25°C, respectively. Bees in the control group tended to either maintain brood care or transition to other active behaviours (i.e., being active on the brood to exhibiting activity off the brood), with a maximum of 1.5% transitioning to inactivity when at both temperatures.

#### 2.2.2 Workers on the brood

The raw number of bees on the brood increased for the control groups at both ambient temperatures (Fig. 3a, Table S10), but due to population growth, the proportion of bees on the brood remained the same. In the pesticide-exposed groups, the proportion of bees on the brood declined over time (*pesticide x time*: t = -6.08, p < .001; Fig. 3b, Table S11) due to a lack of population growth coupled with declines in brood attendance. There was a significantly higher proportion of workers on brood at 19°C and a lower proportion of workers on brood in the pesticide-exposed groups (*temperature x pesticide*: t (177) = -2.13, p = 0.035). Whilst the proportion of bees on brood in the control cohorts remained relatively consistent over the 5 days, there was a steep decline in the proportion of pesticide-exposed bees, with a significant divergence between the control and pesticide groups which began from day 2 of the experiment when at 25°C (W = 2, p = 0.006), and from day 4 at 19°C (W = 14.5, p = 0.008).

**Fig. 3.**
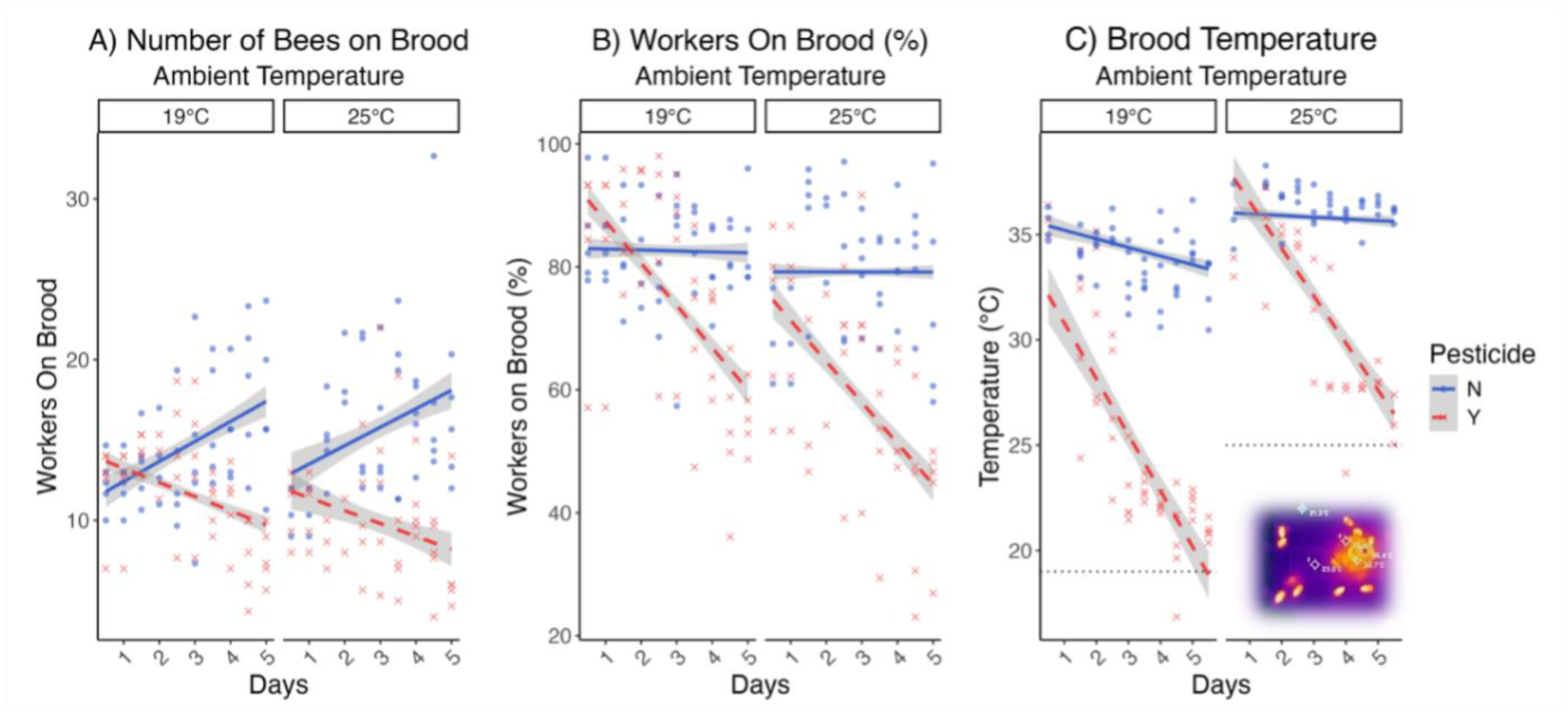
Chronic effects of pesticide exposure on bumblebees leads to decreases in brood attendance behaviour with associated reductions in brood temperature (experiment 2). A) Raw number and B) percentage of workers present on the brood in pesticide-exposed (red crosses, dashed line) and control (blue circles, solid line) cohorts at the two ambient temperatures. C) Surface temperature of the brood in pesticide-exposed and control cohorts. Lines represent model estimates with grey ribbons showing 95% CIs. The dashed lines in plot b show the set ambient temperature (n=20 cohorts, total data points = 185). Inset is a thermal image of an experimental microcolony with nest and workers.

#### 2.2.3 Brood temperature

We found a chronic effect of pesticide exposure that was greater at the lower temperature (19°C) as evidenced by the three-way *pesticide* x *temperature* x *time* interaction (t = 2.35, p = 0.021; Fig. 3c, Table S12). Brood thermal profiles of the control group remained consistent over the 5 days, with a mean brood hotspot temperature in the control colonies of 33.7°C (var= 2.09) at 19°C ambient and 36.2°C (var= 0.60) at 25°C ambient. In contrast, pesticide-exposed colonies experienced significantly lower temperatures compared to the control at both the 25°C ambient temperature (W = 70.5, p < 0.001) and 19°C ambient temperature (W = 120, p < 0.001). The thermal profiles of the pesticide-exposed groups varied more, with a mean brood hotspot temperature of 24.3°C (var=20.08) at 19°C ambient, and 30 °C (var=13.73) at 25°C ambient. By the end of the experiment, brood temperatures of pesticide-exposed cohorts were only ∼1-3°C above ambient in both ambient temperature treatments, whilst control colonies were able to maintain brood at around 30-37 °C.

#### 2.2.4 Testing the relationship between worker brood attendance and brood temperature

The temporally repeated video recordings and brood thermal imaging were conducted in tandem, enabling us to examine how the proportion of bees on brood was associated with brood temperatures. We found a positive relationship driven primarily by the pesticide-exposed cohorts, particularly at the lower temperature (*Pesticide x Temperature x Average bees on brood*: t = 2.18, p = 0.031; Table S13). Whilst generally having more pesticide-exposed bees on the brood appeared to increase brood temperature, this was not the case for control colonies in which brood temperatures remained high regardless of the number of bees on the brood (Fig. 4).

**Fig. 4:**
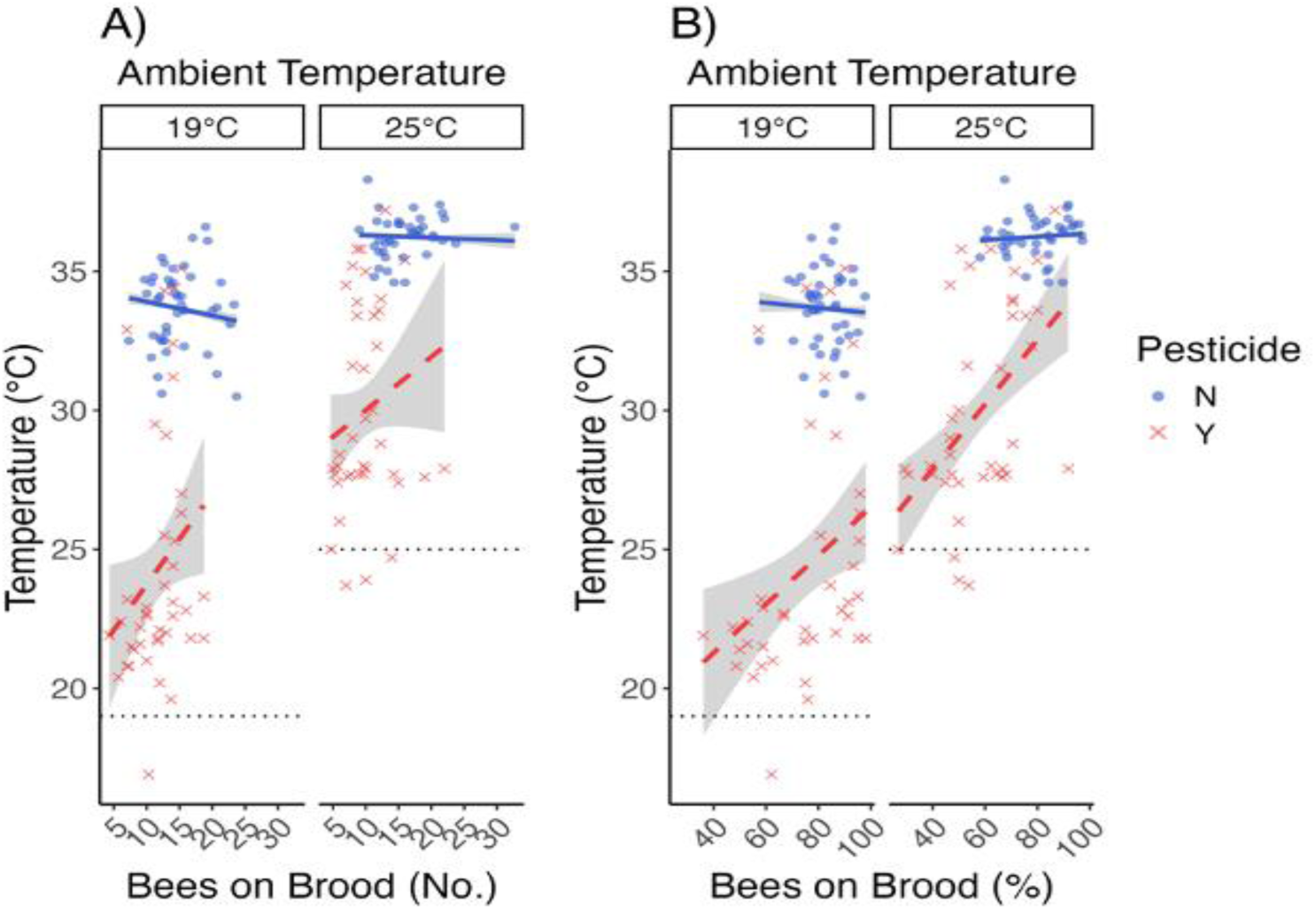
Relationship between brood attendance (x axis) and nest temperature (y axis) supports pesticide exposure leading to ineffective brood thermoregulation (experiment 2). Even when the A) proportion and B) raw number of adult bees on the brood in pesticide-exposed cohorts (red crosses, dashed line) overlaps with control (blue circles, solid line) the surface brood temperature in pesticide-exposed cohorts was significantly cooler on average. Lines represent linear model estimates, with grey ribbons showing 95% CIs. Repeat measures of 20 cohorts, total data points = 185.

#### 2.2.5 Adult population growth

The total number of bees present in colonies at the end of the five-day experiment (sum of surviving original adults and emerged bees whilst accounting for any mortalities; Table S14) was significantly lower in pesticide-exposed cohorts by a mean of 27.7% at 19°C and 39.1% at 25°C relative to control (W = 10.5, p = 0.016; Fig. 5; Table S15). Considering only surviving adults, pesticide-exposed cohorts similarly had lower final populations compared with control groups (W = 5, p = 0.003), with a population decrease at 19°C (mean = 14.4 final adults), and no population change on average at 25°C (mean = 15.1). In contrast, control cohorts grew by a mean of 46.6% at 19°C (mean = 21.9) and 75% at 25°C (mean = 26.3). Data collected on visible cocoon hatching suggests the lower population growth in pesticide-exposed cohorts was due to limited adult emergence. This is because we visibly observed no eclosed (empty) pupal cocoons in the pesticide-exposed cohorts at either temperature, compared to a mean of 3.60 and 5.75 visibly eclosed cocoons in the control at 19°C and 25°C, respectively. Wilcoxon tests showed a significant difference in eclosion associated with pesticide exposure (W = 5, p = 0.002), but not temperature (W = 28, p = 0.50).

**Fig. 5.**
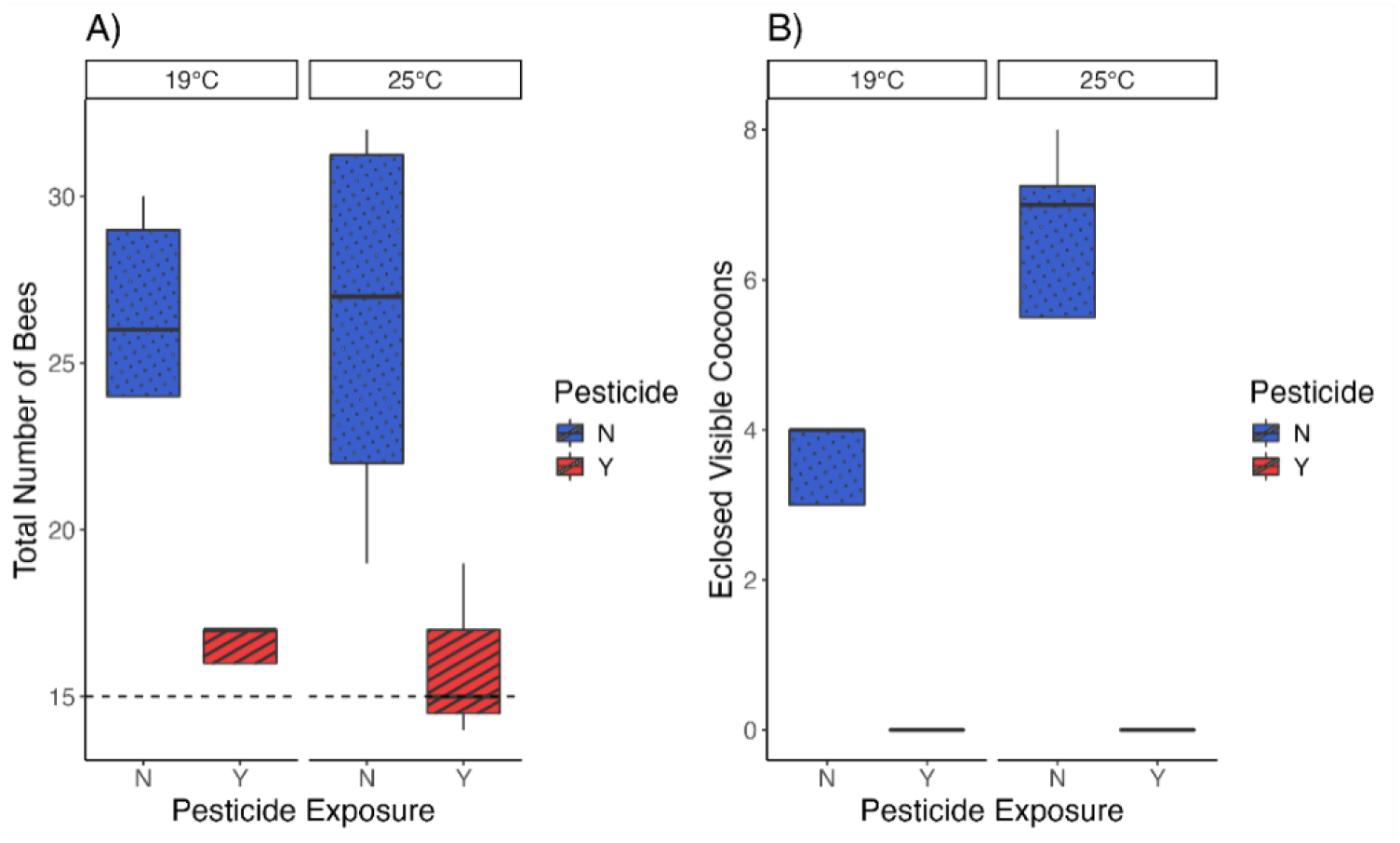
Pesticide-exposed microcolonies having a significantly lower adult bumblebee population at the end of the assay (experiment 2). A) Boxplots showing the total number of adult bees present at the end of the five-day trial, reflecting a dramatically lower net increase in pesticide-exposed cohorts (red striped box) relative to control (blue dotted box; p < 0.01). The dashed line represents the standardised number of adult workers that started the trial in each cohort. B) Boxplots showing the number of cocoons that had visibly hatched from video images by the end of the five-day trial. The solid lines represent the median, the box the interquartile range, and the whiskers the 95% CIs (n = 20 cohorts across treatments).

### 2.3 Experiment 3: Temperature dependent developmental rates of control and pesticide-exposed pupae

Over the 15 days of *in vitro* monitoring, the proportion of pupa that reached our defined final stage of development did not significantly differ between treatment cohorts (control 25°C = 0.61; pesticide 25°C = 0.81; control 34°C = 0.75; pesticide 34°C = 0.65; Pearson’s χ² = 4.95, df = 4, p = 0.29; Table S16). However, the rate at which they reached this stage did. The lower temperature imposed a strong limiting factor on development rate (LM: F(4, 72) = 80.44, p < 0.001; Table S16), with pupae reared at 25°C taking an estimated 5.24 days longer relative to 34°C (when accounting for pupal age at the start of the assay; β = -0.84, p < 0.001). We found no evidence that direct effects of pesticide exposure during development was causing the delayed pupal development rates, as we found no independent effect or interactive effect with temperature (β ≤ 0.93, p ≥ 0.149).

**Fig. 6.**
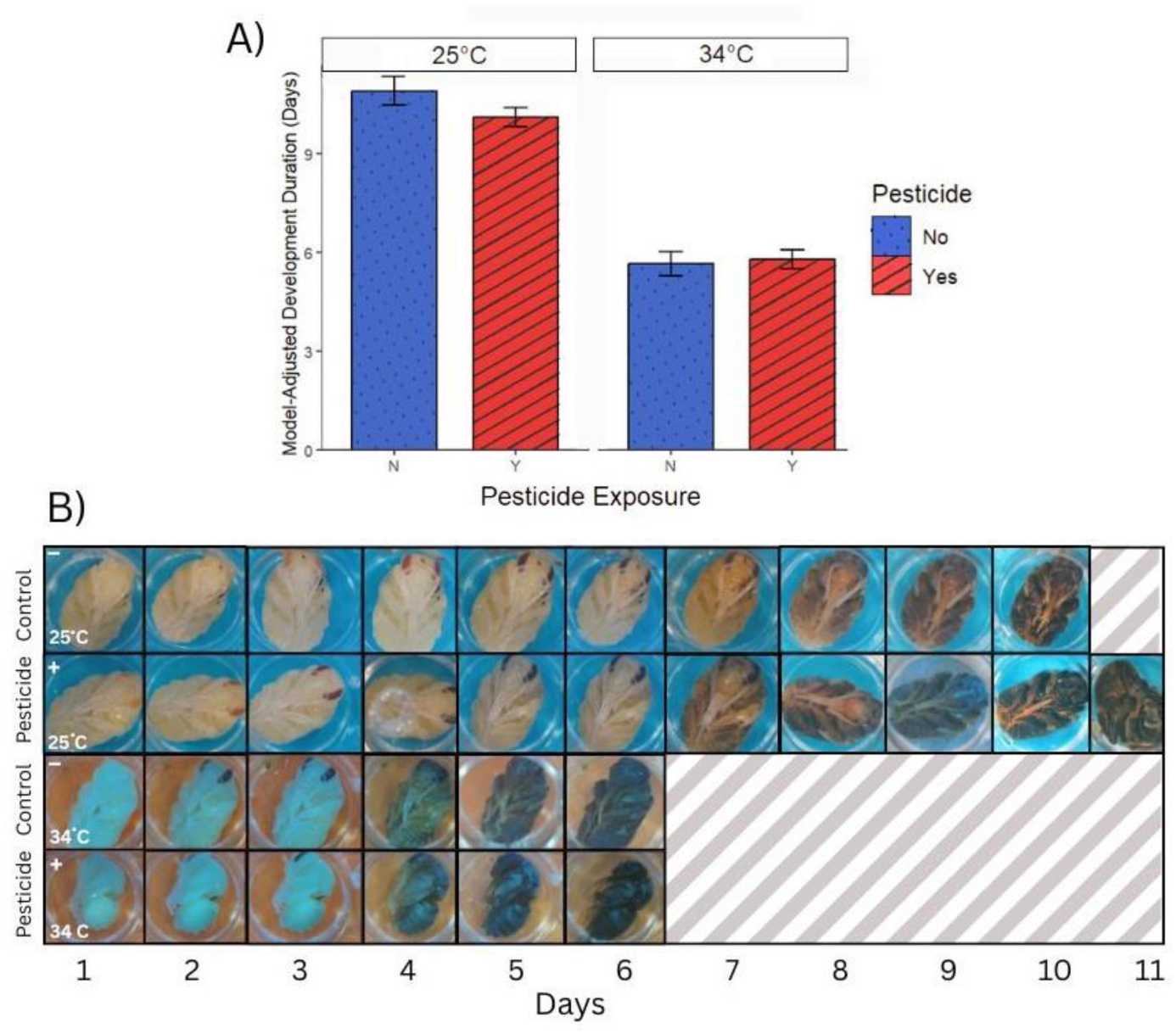
Reduced pupal development rate is driven by lowered brood temperature (an indirect effect of pesticide exposure on nest thermoregulation) and not the direct impact of pesticide toxicity on the individual pupae (experiment 3). A) Bars representing linear model marginal mean estimates with standard error (accounting for initial developmental stage) showing significant effect of temperature (total = 77 pupae). B) Time series images of representative individuals taken daily throughout the pupation period, showing the typical morphological changes from initial eye pigmentation to fully melanised cuticle before emergence.

## 3. DISCUSSION

We show that pesticide exposure (a neonicotinoid in this case) prevents adult bumblebees from maintaining a suitable [above ambient] body temperature whilst decreasing behavioural activity, including brood attendance. This dual impairment occurred quickly after the onset of exposure, and led to brood temperature gradually, but distinctively, dropping. This effect was exacerbated at the experimental lower temperatures (15-20°C), reinforcing the notion that neonicotinoids impair non target insect thermogenesis (in this case the ability of a bee to warm up (32, 33)). Indeed, our analyses further showed that the efficiency to which workers can keep the brood warm was undermined by neonicotinoid exposure. Concerningly this was associated with little to no growth of the colony adult population and our in vitro rearing assay supports that this is due to the indirect effects of exposure on brood thermoregulation rather than due to direct physiological effects from the pesticide.

### 3.1 Pesticide-exposed bumblebees could not maintain body temperature

Experiment 1 showed that bumblebees (at least *B. terrestris*) have an inherent behaviour of heating themselves above the ambient temperature (even when not in flight (42)), and that neonicotinoid exposure places this adaptation at risk. Generally, we found control bumblebee adults consistently held their body temperature over six hours between an average of 29 °C (at 15 °C ambient) and 32 °C (at 25°C ambient), with <3% variation from trial start to end. This range of non-flight body temperatures aligns with reported optimal brood rearing temperatures (43), supporting individual thermoregulation to be an evolved response for enabling brood thermoregulation. But we also show that neonicotinoid exposure can leave adults unable to maintain suitable body temperatures, with a 21.3%, 6.6%, and 7.3% average reduction in maximal thorax surface temperature over the six hours when at 15, 20 and 25 °C, respectively.

Whilst we suspect pesticide effects on body temperature to be an issue for other insect taxa, impacts on generating and maintaining body temperature in bumblebees is problematic given they are heterotherms and as cold adapted specialists are frequently required to heat their body for important tasks (42, 44). Our findings can explain why previous results have shown temperature dependent effects of neonicotinoid toxicity on bumblebee locomotive behaviours (21) and survival (23) and expand on previous work showing that imidacloprid exposed bumble bees (32), honey bees (23), and solitary bees (45) struggle to survive and recover from cold temperatures. The worsening effects of lower ambient temperatures for pesticide-exposed bees over time suggests that with projected increases in the frequency and intensity of extreme low temperature events (46), alongside the ubiquity to which bees are exposed to pesticides (39), individuals could be at risk of becoming less active at key phases in the season.

Past studies have shown neonicotinoid exposure to be associated with reduced appetite in bees (21, 47), potentially leaving bees with less energy for thermogenesis/endothermy (28). Our results showed, however, that appetite responses are non-uniform. Neonicotinoid exposed bees consumed more food relative to control bees under the lower temperature (15 °C) but in contrast consumed less at the high (25 °C). Due to the declines in thorax temperature in pesticide-exposed bees at the lowest ambient temperature, it appears the elevated food consumption did not enable pesticide-exposed bees to compensate for the pesticide effects. This points towards a direct effect of the neonicotinoid on molecular and neuronal pathways preventing bumblebees from being physically capable of controlling their own body temperature. Previous neonicotinoid exposure experiments in bumblebees have shown the expression of genes involved in energetic metabolic pathways to be affected (27, 48, 49), and that mitochondrial dysfunction may be symptomatic of exposure (29, 30). It thus seems that internal pathways involved in thermogenesis and homeostasis are being affected, as suggested in honeybees (33).

Intriguingly, at our highest tested ambient temperature (25°C), neonicotinoid exposed bees experienced an acute effect of higher thorax temperature relative to controls in the first 2-3 hours, followed by a continual drop until the end of the six-hour trial. High ambient temperatures are likely to be associated with increased rates of pesticide breakdown in insects (50), though this depends on the mode of action (51). The drop in thorax temperature by pesticide bees to then converge with control bees may reflect an element of recovery or acclimation from this impairment, but with body temperature continuing to decline, it could also be attributed to a total collapse in heat generation. That said, previous work with honeybees has reported imidacloprid exposure to increase survivable thermal maxima (52), which if true could suggest that our observation was a result of a reduced need to cool themselves.

### 3.2 Individual impairment from pesticide exposure translates to reduced colony-level thermoregulation leading to delayed brood development and lower adult population growth

Whilst control cohorts maintained an ∼34-36 °C brood temperature over the trial (aligning with that expected; (43)), neonicotinoid cohorts at 19 °C and 25°C exhibited a gradual but distinct 41.7% and 20.0% average reduction in brood temperature, by the end of the trial, respectively. For control cohorts, there was no relationship between the number of workers attending brood and brood temperature, indicating that just a relatively small number of workers are needed to maintain thermal homeostasis. Yet, we found a positive relationship for pesticide-exposed cohorts indicating the need to compensate for neonicotinoid induced reductions in individual body temperature. However, even where a high proportion of workers were observed on the brood, brood temperatures still rarely reached that of controls. To compound this issue, once around day 3 had passed, worker collective behaviours in cohorts showed a distinct change, with a significant drop in active workers and brood attendance. Such impacts on collective behaviours echoes the conclusions by Crall and colleagues (10) who reported neonicotinoid impacts on social interaction networks within bumblebee colonies associated with a drop in nighttime colony temperatures. For our studied bumblebees, the consequences of an inability to rear a large enough adult population are that brood care and foraging activities will not be able to keep up with the energetic needs of overlapping adult generations (53). For instance, in *Bombus huntii*, experimental removal of incubating bees can increase incubation times of remaining brood carers, rather than task switching of other bees to incubation (Gardner et al., 2007). Though some thermal buffering can be achieved by having underground nests, trade-offs between trying to counter brood getting cold with other important colony functions (e.g. foraging to meet metabolically energetic demands) could follow Allee type patterns, reinforcing negative feedback loops and leading to colony collapse (5, 54).

Using a combined colony level exposure followed by *in vitro* assay (experiment 3), we looked to test whether delays in developing pupae was indeed down to the indirect effects of pesticide affecting nest temperatures, or because the direct toxic effects of the pesticide itself, or both. By rearing the control and exposed pupae in vitro at temperatures measured in experiment 2 for end of trial control nests (34 degrees) and neonicotinoid nests (25 degrees), we found evidence for the indirect hypothesis. Indeed, brood temperature should be important for successful larval and pupal development of many holometabolous insects, particularly in bumblebees (24–26, 55).

Our findings contribute a potential mechanism explaining why lower productivity in bumblebee colonies has been found near pesticide treated agricultural fields (14, 56) and lower offspring hatching in solitary bee nests in which parents have been foraging on neonicotinoid treated plants (57). It can further explain why controlled neonicotinoid exposure of bumblebee colonies in the field by Arce and colleagues (12) found lower numbers of reared gynes and males despite foraging patterns appearing little affected (also see (17)). It should be noted, however, that in the study presented here, we used cohorts of 15 workers which is smaller compared to typical field colony sizes nearer the end of the season (>100 workers). Larger bumblebee colonies have been suggested to better buffer impacts of imidacloprid on thermoregulation and population growth (58), yet with pesticides heavily applied at the start of the season, colony sizes are naturally small at that time.

### 3.4 Conclusion: implications for bumblebees and other insect pollinators under future landscapes

Colonial living in social insect is considered an adaptation for buffering environmental pressures, such as temperature extremes (59). Our result, however, suggests pesticide exposure undermines this strategy. We show explicitly that pesticide effects on thermogenesis and nest thermal homeostasis translates to impacts on brood development, and yet this is unlikely to be exclusive to bumblebees as other insects require suitable body temperatures for foraging and suitable nest temperatures for rearing offspring (60, 61). The multi stressor nature of modern landscapes suggests that care should be given when assessing the context dependency of pesticide risks (21, 62). Furthermore, in environments where food resources are heterogeneous and there is competition for resources, the additional impairments that pesticides have on locomotion (21, 47, 63) and cognition (47, 64) could exacerbate the situation. Further research is needed to understand multi stressor effects on beneficial insect fitness, particularly temperature and pesticide combinations, to inform targeted mitigative action and to inform risk assessments and environmental policy. Our findings put forward a key mechanism for how relatively recent global change factors may have driven population declines of key beneficial insects.

## 4. MATERIALS & METHODS

### 4.1 Bumblebee colonies, and controlling pesticide & temperature exposure

We obtained five, eight, and five *Bombus terrestris audax* colonies for experiment 1, 2 and 3, respectively, to act as parental donors for our assays from the commercial supplier Biobest (distributor Agralan Ltd). Parental donor colonies (all experiments), individual pots (experiment 1) and microcolonies (experiment 2) were held in controlled environment (CE) rooms at 60% relative humidity. For all experiments, the *ad libitum* nectar substitute that bees were provisioned was a 40% w/w sucrose/water solution, with the pesticide treatment group being spiked with 10 μg/L imidacloprid representing a concentration found in nectar and pollen of flowers in the field (65). For experiment 1, focusing on individual relative surface body temperatures, we tested bee responses at ambient temperatures of 15, 20 and 25 °C. For experiment 2 on whole cohort responses, we tested under 19 and 25 °C to mimic mean Spring-Summer temperatures recorded at Silwood Park (Ascot, UK) from 2010-2019 (66) and within the range tested in experiment 1. For experiment 3, we reared pupae *in vitro* at 25 °C and 34 °C to simulate approximate surface brood temperatures that neonicotinoid exposed and control microcolonies (respectively) showed near the end of experiment 2 (see Results 3.2.3).

### 4.2 Thermal imaging (experiments 1 and 2)

A FLIR ONE Pro camera (FLIR Systems) captured thermal images (emissivity set at 0.95) of test subjects (Fig. 1). Pilot work testing the thermal imaging showed that use of black electrical tape (3M Scotch super 33+ PVC tape with known emissivity of ε = 0.95 ±. 0.05) enabled us to calibrate camera readings with high accuracy (see Supplementary Methods). FLIR images were processed using a hotspot tool (FLIR Tools v.5.13.18031.2002) to return relative thermal readings for each subject. When imaging subjects, we found that 93% of images taken by the camera read the tape temperature within 0.2 °C of the known and tightly controlled room temperature (21), and we further excluded any images above this margin of error. For this comparative treatment experiment, this provided a suitable standardised process to thermally profile between adults and between brood to gain relative comparisons of thermoregulation at the individual and colony-level. For experiment 1, hotspots always represented a thoracic pixel of a bee. For experiment 2, if hotspots were not automatically assigned to the brood by the FLIR tools package, polygons were manually assigned to represent the brood area with the hotspot then extracted from this portion of the image. We validated that the thermal hotspot could be used as a proxy of wider brood relative thermal profiles by showing that the hotspot has a strong correlation with the mean temperature of three randomly selected points across the brood (R^2^ = 0.85; Fig. S2, Table S1).

### 4.3 Experiment 1: Temperature dependent effects of pesticide exposure on individual thorax temperature profiles

On arrival, parental donor colonies were allowed to acclimate to lab conditions for five days before experimental assays started. These parental colonies were kept in a red-lit CE room set at 25 °C and provided with *ad libitum* plain nectar substitute and four grams of honeybee collected pollen daily (BioBest). For the trial we took individual workers from their parental colony (Table S2) and placed each in a separate container (120 mL) for the duration of the experiment. In total, we randomly selected 127 workers (24-28 per parental colony, see Supplementary methods for protocol).

For each of the three temperature treatments (n=37-43 workers per temperature), half the workers were fed either the untreated or pesticide spiked nectar substitute. This was done via an Eppendorf tube in which a small hole was made at the bottom (using tip of a soldering iron) and then the tube being inserted in the pot side wall just above the floor. We measured the mass of nectar substitute consumed finding no significant difference between the control and pesticide-exposed groups (t-test = -1.24, df = 71.7, p = 0.22). Beginning when the food was provisioned, thermal images (taken once the pot cover was removed) were captured at 60 minutes, 120 minutes, and then at 30-minute intervals until six hours had elapsed, resulting in a total of ten images. Six hours was selected to ensure metabolism of imidacloprid (67) whilst maintaining a feasible recording period. For each thermal image, six pots were configured in a way that allowed thermal profiling of six workers under a single field of view. At the end of the six-hour trial, workers were frozen at -20 °C, with individual dry mass calculated post-trial by drying in 80 °C ovens for 72 hrs and weighed.

### 4.4 Experiment 2: Temperature dependent pesticide effects on worker activity, brood care behaviour and cohort brood temperature

Within two days of arrival, from each parental donor colony we created 2-3 cohorts with each cohort comprising fifteen randomly selected worker bees (twenty cohorts in total; Table S3). Each cohort was provisioned 10-16g of the parental colony’s brood with a balanced composition of comb type (same approximate ratio of eggs, larvae, pupae, and nectar pots), with no significant difference in brood mass between treatments (see Table S4). The resultant ‘microbroods’ were placed on a 45mm diameter plastic petri dish in the middle of a ventilated high-density polypropylene microcolony box with the selected workers. The cohorts (microcolonies) made from each parental colony were evenly split between our two experimental temperatures (19 or 25 °C) and allowed to acclimate for two days with *ad libitum* provision of nectar substitute via gravity feeders and 2.5 g of honeybee collected pollen inside a 45mm petri dish provisioned on days 0, 2 and 4. Any workers found dead prior to the start of the assay were removed and replaced with a randomly chosen worker from the respective parental colony to maintain the worker population at fifteen at the start of the trial.

After acclimation, half of the cohorts in each CE room were assigned to control and the other half to pesticide treatment. Both groups were supplied with 2.5g of pollen on days 1, 3, and 5 of the five-day experiment. Nectar consumption was measured by weighing the tubes daily, and pollen consumption by weighing the pollen dish at the end of the trial. Mortality was visually recorded each day with dead workers removed and frozen at -20 °C. At the end of the experiment, all cohorts were frozen (-20 °C) and later censused with final mass and composition of the brood recorded. Given that the duration to develop from a first day pupa to an adult bumblebee is more than five days, the emergence of any bees during our five-day experiment will be indicative of pupal development. Crucially pupae do not require any further food provision from adult workers, but their development rate and likelihood of eclosion should be dependent on temperature (43).

Thermal images were taken in triplicate twice a day for each cohort (i.e., total of six images per day per microcolony). Top-down video recordings of within cohort behaviour was conducted for three minutes in the morning and evening prior to thermal imaging using either a Cannon (LEGRIA HFR606) or Panasonic (HC_V160) camera. Behavioural data was obtained by manually scoring behaviours for each cohort individual at the 60^th^, 120^th^ and 180^th^ second in the footage for i) the ratio of bees in contact with brood and ii) the type of behaviour each bee was exhibiting, categorised as either active on the brood, active off the brood, or inactive based off an ethogram (Table S5).

### 4.5 Experiment 3: Temperature dependent developmental rates of control and pesticide-exposed pupae

On arrival parental donor colonies were maintained at 25°C and allowed to acclimate for 1 week with pollen provided *ad libitum* and untreated sucrose solution *ad libitum* via gravity feeders. Once the week was up, three colonies were randomly assigned as the pesticide exposure treatment and two colonies as control. We continued this exposure for seven days because this is the average time taken to develop from an egg to a 1-day old pupa, and thus the early-stage pupae in the colonies at this end point must have developed during the parental colony exposure treatment period. After this week of exposure, brood cells (pupal cases) were collected from each colony (n = 20-30 per treatment). Early-stage pupae could be distinguished based on the texture and colour of the pupal case (lighter colour indicating younger). Then, once we had removed the pupae from the case, we could more accurately determine age (development stage). Pupae were aseptically removed from the pupal case using sterilized forceps. Individuals were selected at either the final larval stage or early pupal stage, the latter defined by the presence of white eyes, indicating the onset of pupation (68). Pupae were placed individually into sterile 48 well plates (1ml) wells, with lids modified by drilling 2mm holes to allow for airflow. These plates were then placed into sealed plastic containers (Tupperware, 10 x 5 x 5 inches), containing saturated NaCl solution to maintain a stable relative humidity of 75%, following standard bee rearing protocols (69, 70).

The isolated pupae from each parental colony were then divided equally and reared at either 25°C or 34°C (representing nest temperature results from pesticide and control colonies from experiment 2, respectively) to assess if pupal development is affected by temperature, the prior pesticide exposure environment, or an interaction between the two. Of the 111 pupae that started the *in vitro* rearing assay, five were removed within 24 hours due to signs of damage during the extraction and transfer process, with the probability of exclusion not significantly differing between the four treatments (*p* = 0.89, χ² = 0.58, df = 3). This resulted in the following treatments and sample sizes: 25 °C control n = 21; 25 °C pesticide-exposed n = 31; 34 °C control n = 33; 34 °C pesticide-exposed n = 21 (Table S16). Pupal development was monitored daily using a digital stereomicroscope fitted with a high-resolution camera (Andonstar). Individual images were taken at the same time each day to ensure temporal consistency with key developmental stages timestamped using photographic records, with reference to described morphological criteria (68). For this study, we classified the end of pupation as the point at which individuals displayed a fully melanized cuticle and spontaneous leg twitching, immediately after which each respective worker offspring was removed and frozen to avoid bees developing into mobile adults and escaping.

### 4.6 Statistical analyses

Analyses were run using R statistical software via RStudio using either Version 2023.06.2+561 or 2025.05.0 Build 496 “Mariposa Orchid” (71) with graphical plots made using ggplot2 (72). For model construction, we first considered maximal models with all explanatory variables (using appropriate model functions according to data distributions) before dropping components and using AIC to select the best fit model (73). Best fit models were then used to produce relationship estimates (with 95% CIs), with p-values computed using a Wald t-distribution approximation, producing a Nagelkerke’s R value (74).

#### 4.6.1 Experiment 1

Food consumption was square root transformed to improve normality, and dry mass log transformed given the relationship between thermogenesis and body mass likely follows a power law (75). We then calculated consumption per unit body size (*mass*/*food consumption)* to collapse body size and food consumption into a single variable. The best fit model for predicting thorax temperature was a generalised linear mixed model (GLMER) fit with an identity link function. The model contained two-way interactions of *pesticide x temperature*, *pesticide x time*, and *mass*/*food consumption x pesticide.* The model included time as a random effect that varied with bee identity to account for the time series data structure (i.e., repeat measures of individuals; see Tables S6 and S7 for model structures) and had a total explanatory power of 65% of the thorax temperature variation.

#### 4.6.2 Experiment 2

Brood temperature data were averaged across the three thermal images captured per time step, and log transformed to account for mass impacts on temperature and improve normality. The best fit model was a fixed effects GLM using a square root link function and structured as a three-way interaction between *pesticide* x *temperature* x *time*, and considering additive effects of *pre-trial brood mass* and *cohort ID*. The model’s total explanatory power was 88% of brood temperature variation.

The behavioural data of the cohorts was first analysed as worker attendance to the brood (proportion of bees on the brood). The best fit model for brood attendance was a fixed effects linear model (LM) with a three-way interaction of *pesticide* x *temperature* x *time*, with an additive effect of *parental colony of origin*. The model’s total explanatory power was 58% of the data variation. Behavioural data was then further analysed by calculating the transitions between behaviour types between day 1 and 5 of the experiment (active on the brood, active off the brood, or inactive) to assess trends in behaviour transitions between trial groups. We explored how behavioural data could predict thermal profile data by using a GLM that included a three-way interaction between *pesticide* x *temperature* x *time* and the proportion of bees on brood as an additive effect. The model’s total explanatory power was 87% of brood temperature variation.

Data on brood mass and number of bees at the end of the trial were analysed using Wilcoxon tests to assess significant differences in final cohort population (starting adult bees and emerged adult bees minus mortality) and pupal emergence (visible eclosed pupal cells) between the trial groups.

#### 4.6.3 Experiment 3

To test the interactive effect of *temperature x pesticide* on pupal development rate we fitted a linear model (LM) while controlling for variation in initial developmental stage (*start age*). Because individuals entered the experiment at different developmental stages, an estimated *start age* was assigned based on visible eye and cuticle pigmentation and included as a covariate in the model. This correction ensured that comparisons reflected true differences in developmental rate rather than differences in starting stage. Post hoc pairwise comparisons of estimated marginal means were performed using the emmeans package (76).

## Acknowledgments

Thanks to Ana Ramos Rodrigues for helping with husbandry and setups, Paul Beasley and Martin Selby for technical support, and the Gill and Graystock groups for feedback on manuscript drafts.

## SUPPLEMENTARY INFORMATION

### Supplementary Methods

#### Experiment 1 – parental colony husbandry and selecting bees for the trial

On arrival the sugar solution reservoirs that came with the parental colonies were immediately removed. We then provisioned ad libitum plain 40% sucrose solution along with a daily amount of 4g of pollen.

For the experimental trial (6 hours), each individual bee did not have access to pollen inside their respective holding pot. Bees were selected for the experiment from across the parental colonies. Trial replicates 1 and 2 had 12 bees, but sample size was later reduced to eight bees due to imaging asynchrony. Experiments were then run with eight workers a time (four control, four imidacloprid-exposed per temperature room). Trial replicate 16 had five bees tested to correct for missing data in the initial trial replicates. Bees were selected uniformly from the parental colonies apart from an early preference for colony 6 due to its large size and to allow the other four parental colonies to grow (Table S2).

#### Experiment 1 and 2 - thermal imaging camera accuracy

Trial data was collected with the FLIR ONE Pro camera (FLIR Systems) to test whether the camera settings interfered with temperature measurements. The camera was held in a clamp 0.5m above the black tape used for calibration. Readings were taken of the temperature of the tape every 10 seconds over a period of 10 minutes from camera switch-on, to observe if/how readings changed over time. Temperature readings did converge, but at 25 °C the camera consistently recorded the wrong ambient temperature. This revealed that the camera’s autocalibration was causing error in true temperature readings due to changes in shutter speed associated with warming. We therefore tested accuracy when the autocalibration was turned off and found that temperature readings were then consistent with the ambient temperature. For our trials, therefore, all autocalibration settings were turned off.

#### Experiment 2 - measuring microcolony brood temperature

To check that the hotspot of brood is representative of the general brood temperature three random points were selected on the brood (n = 87), averaged, and compared with the hotspot value. A simple linear model using *hotspot x pesticide exposure* as explanatory variables for the average temperature of the other datapoints showed a tight relationship with R^2^ = 0.879 (Fig. S1; Table S1).

#### Experiment 1 and 2 – statistical model selections

Individual thermal data, microcolony thermal data and proportion of workers on brood, initial maximal mixed effects (lmer or glmer) and fixed effects (lm or glm) models (package lme4; Bates et al., 2015). Complete global model components are shown in Table S6. Global models were simplified through iterative comparisons using analysis of variance (ANOVA) and the output Akaike Criterion Index (AIC) to assess nested model fit (Akaike, 1974) and to assess what combination of traits adequately reduces model complexity whilst retaining explanatory power (Mazerolle, 2023). The best fit model output (see Table S7) was summarised using the report package (Makowski et al., 2023), and base R was used to generate predicted model outputs (R Core Team, 2023). The best fit model was used to produce a predicted line of best fit and plotted with a 95% confidence interval along with the raw data points, with p-values computed using a Wald t-distribution approximation, producing a Nagelkerke’s R value (Makowski et al., 2023).

#### Experiment 3 - statistical model selection

To assess the effects of temperature and pesticide exposure on pupal development duration, a linear model (lm) was constructed including temperature, pesticide exposure, their interaction, and start age (estimated developmental stage at the onset of rearing based on Tian and Hines; 2018) as predictors. Model selection was performed using comparisons of nested models via analysis of variance (ANOVA) and evaluation of Akaike Information Criterion (AIC) values (Akaike, 1974; Mazerolle, 2023). The full model (AIC = 268.4 Table S17) showed a better fit than models excluding the interaction term (ΔAIC = +0.24), pesticide (ΔAIC = +41.8), or temperature (ΔAIC = +69.4), supporting the inclusion of the interaction and both main effects.

## Supplementary Figures

**Fig. S1.**
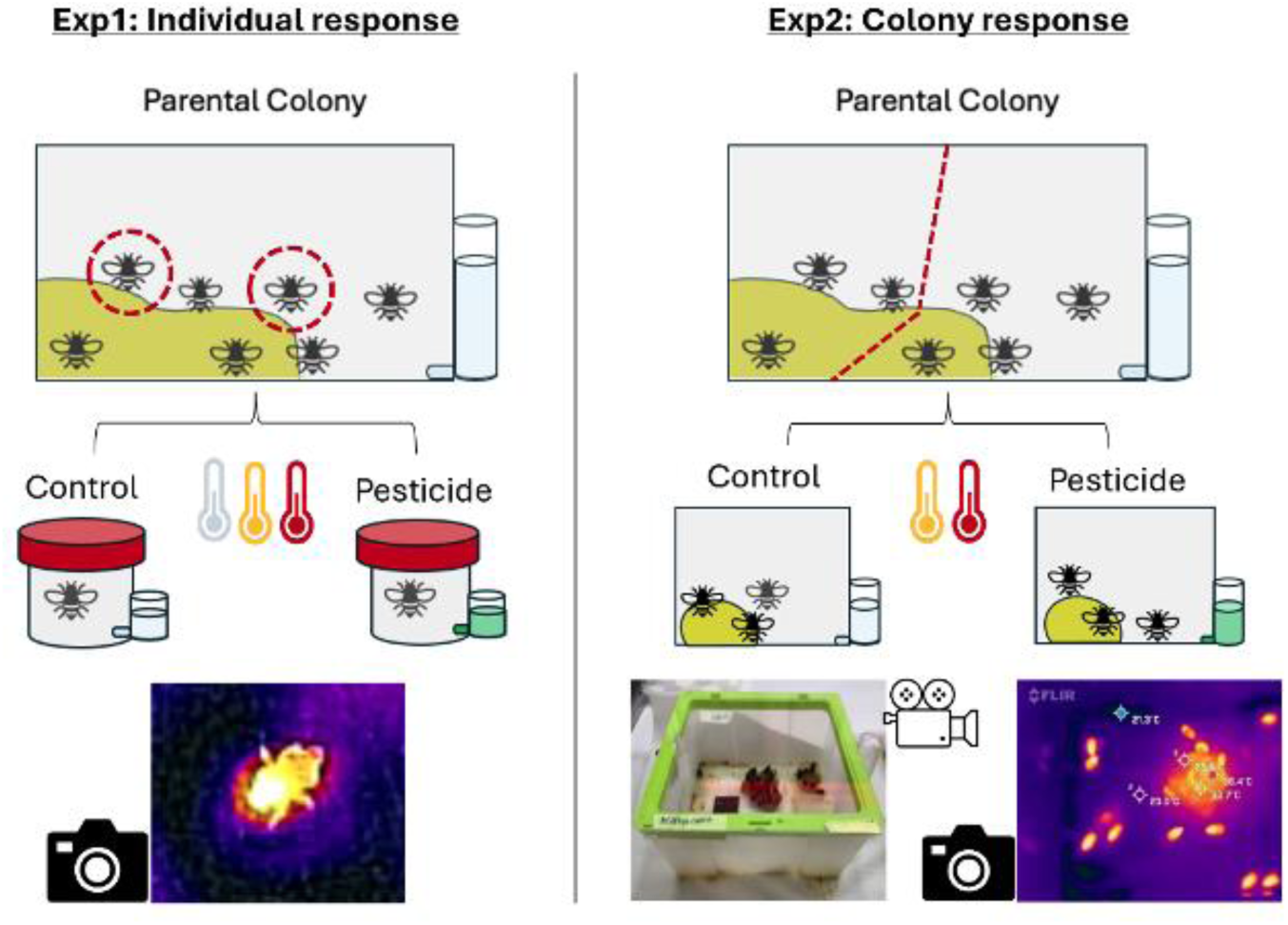
Setups for experiments 1 and 2 showing the temperature and pesticide treatment combinations and response measures. 1: Individual level responses (experiment 1) whereby adult workers were removed from their parental colony and separately held in pots under 15, 20, or 25 °C. For each temperature cohort, half the bees were provisioned ad-libitum control nectar substitute and the remaining half where it was pesticide spiked. Workers were observed for six hours with thermal images repeatedly taken. 2: Colony level responses (experiment 2) in which fifteen adult workers were removed from a parental colony to make a new cohort. The cohorts were placed under 19 or 25 °C, and for each temperature cohort half the cohorts were provisioned ad-libitum control nectar substitute, and the remaining half pesticide spiked. Cohorts were observed for five days (120 hours) during which worker behaviour and activity were repeatedly video recorded alongside thermal imaging of the brood.

**Fig. S2.**
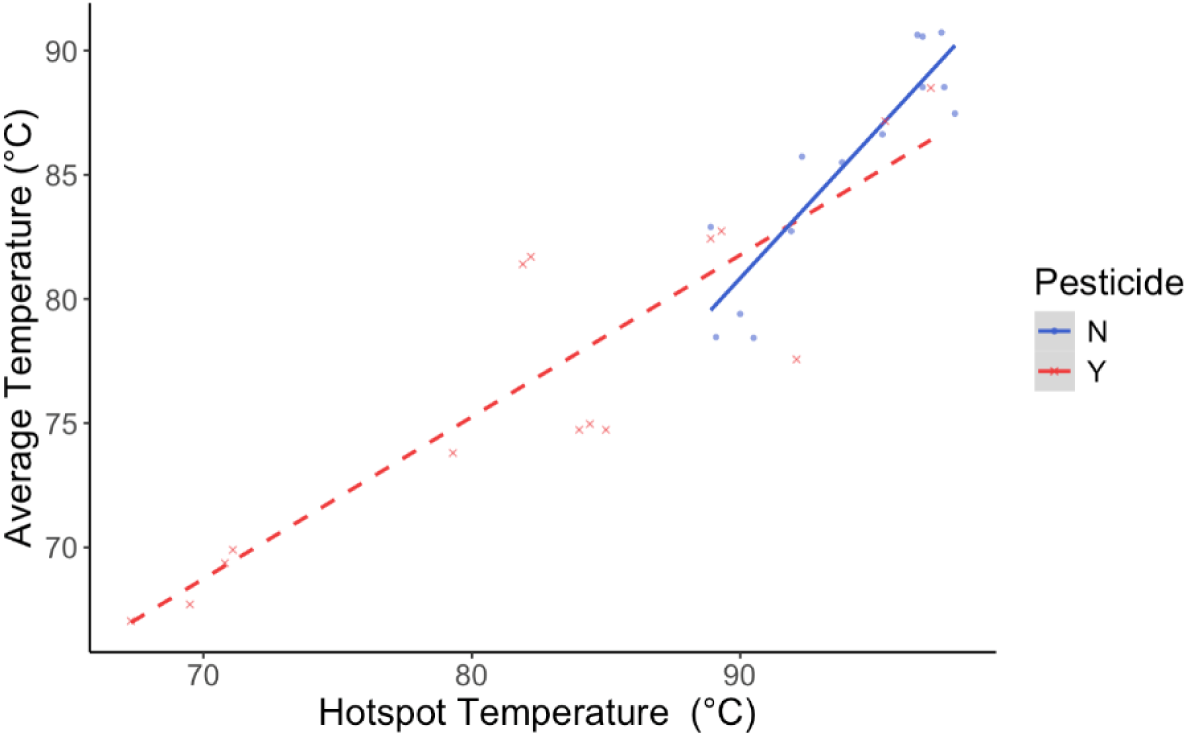
LM (model) estimates of the average colony temperature predicted by hotspot temperature for pesticide-exposed (red crosses and dashed line) and control (blue circles and solid line) microcolonies (n = 29 brood measures).

## Supplementary Tables

**Table S1:** Model output to test if average temperature could predict brood hotspot whilst considering if pesticide treatment altered this relationship.

| *Predictors* | **Average_temp** |  |  |
| --- | --- | --- | --- |
|  | *Estimates* | *CI* | *Statistic* |
| (Intercept) | -24.63 | -66.29 – 17.02 | -1.22 |
| Hotspot brood | 1.17 | 0.73 – 1.62 | 5.45 |
| Pesticide [Y] | 47.71 | 4.14 – 91.29 | 2.26 |
| Hotspot brood × Pesticide [Y] | -0.52 | -0.99 – -0.05 | -2.28 |
| Observations | 29 |  |  |
| R<sup>2</sup> / R<sup>2</sup> adjusted | 0.879 / 0.864 |  |  |

**Table S2:** Number of bees selected from each parental colony for testing in experiment 1 (total = 127).

| Colony | Number of Individuals |
| --- | --- |
| 3 | 25 |
| 4 | 25 |
| 5 | 24 |
| 6 | 28 |
| 7 | 25 |

**Table S3:** Parental colony ID and the number of microcolonies (15 workers per microcolony) created per temperature treatment.

| Parental Colony | Microcolonies | Temperature |
| --- | --- | --- |
| 1 | 3 | 19 |
| 3 | 3 | 19 |
| 11 | 2 | 19 |
| 12 | 2 | 19 |
| 13 | 2 | 25 |
| 14 | 3 | 25 |
| 15 | 3 | 25 |
| 16 | 2 | 25 |

**Table S4:** Wilcoxon Test comparisons of the starting brood mass provisioned for microcolonies between the different sample groups (pesticide and temperature combinations).

| Comparison | W-value | p-value |
| --- | --- | --- |
| Pesticide vs Control | 61 | 0.43 |
| 19°C: Pesticide vs Control | 13 | 1 |
| 25°C: Pesticide vs Control | 6 | 0.22 |

**Table S5:** Ethogram for classifying different worker behaviours from the repeated snapshots of video footage during experiment 2.

|  |  |  |
| --- | --- | --- |
| Inactive |  | Lethargic bee is lying in motionless position, and the bee is not engaged in feeding or nest duty behaviours. |
| Active | Brood | Bee is located on the brood and showing clear signs of defined movement including locomotion, nestmate interactions, nest building, and/or attending and feeding brood. |
|  | Off-brood | Bee is not on the brood but showing clear signs of defined movement including locomotion, feeding, nestmate interaction and/or grooming. |

**Table S6:** Global model structures for the different measured temperature responses across experiment 1 and 2.

| Response Variable | Global Model |  |  |  |
| --- | --- | --- | --- | --- |
|  | Model type | Link function | Components | AIC |
| Individual Thorax Temperature | glmer | Family=<br>Gaussian Link<br>= Identity | Pesticide + Ambient Temperature + Time+<br>Size/food consumption + Bee ID + (Time \|<br>Bee ID) | 3573.9 |
| Brood Temperature<br>(considering<br>bees on brood) | glm | family =<br>Gamma<br>Link = Log | Pesticide * Ambient Temperature * Time +<br>Bees on Brood * Pesticide * Temperature +<br>Starting Brood Mass | 719.49 |
| Brood Temperature (%<br>bees) | glm | family =<br>Gamma<br>Link = Log | Pesticide * Ambient Temperature * Time +<br>Proportion of Workers on Brood * Pesticide<br>* Time + Starting Brood Mass | 724.15 |
| Bees on Brood | lmer |  | Pesticide + Ambient Temperature + Time +<br>(1\|Microcolony) | 1019.24 |
| Proportion of<br>Bees on Brood | lm |  | Pesticide + Ambient Temperature + Time +<br>(1\|Microcolony) | -203.65 |

**Table S7:** Best model structures for the different temperature responses across experiment 1 and 2 based on AIC comparisons showing the variation explained (R^2^).

| Response Variable | Best Model |  |  |  |  |
| --- | --- | --- | --- | --- | --- |
| | Model type | Link function | Components | AIC | $R^2$ |
| Individual Thorax Temperature | glmer | Identity | Pesticide * Ambient Temperature +<br>Pesticide * Time + Size/food consumption* Pesticide + (Time\| Bee ID) | 3547.4 | 0.65 |
| Brood Temperature (number of bees on brood) | glm | family = Gamma<br>Link = Log | Pesticide * Ambient Temperature * Time +<br>Average Bees on Brood * Pesticide * Temperature | 719.34 | 0.88 |
| Brood Temperature (% bees) | glm | family = Gamma<br>Link = Log | Pesticide * Ambient Temperature * Time +<br>Proportion of Workers on Brood * Pesticide * Time | 715.00 | 0.88 |
| Bees on Brood | lmer |  | Pesticide * Ambient Temperature +<br>Pesticide * Time + (1\|Microcolony) | 980.23 | 0.56 |
| Proportion of Bees on Brood | lm |  | Pesticide * Ambient Temperature + Ambient Temperature * Time + Pesticide * Time | -259.83 | 0.50 |

**Table S8:**
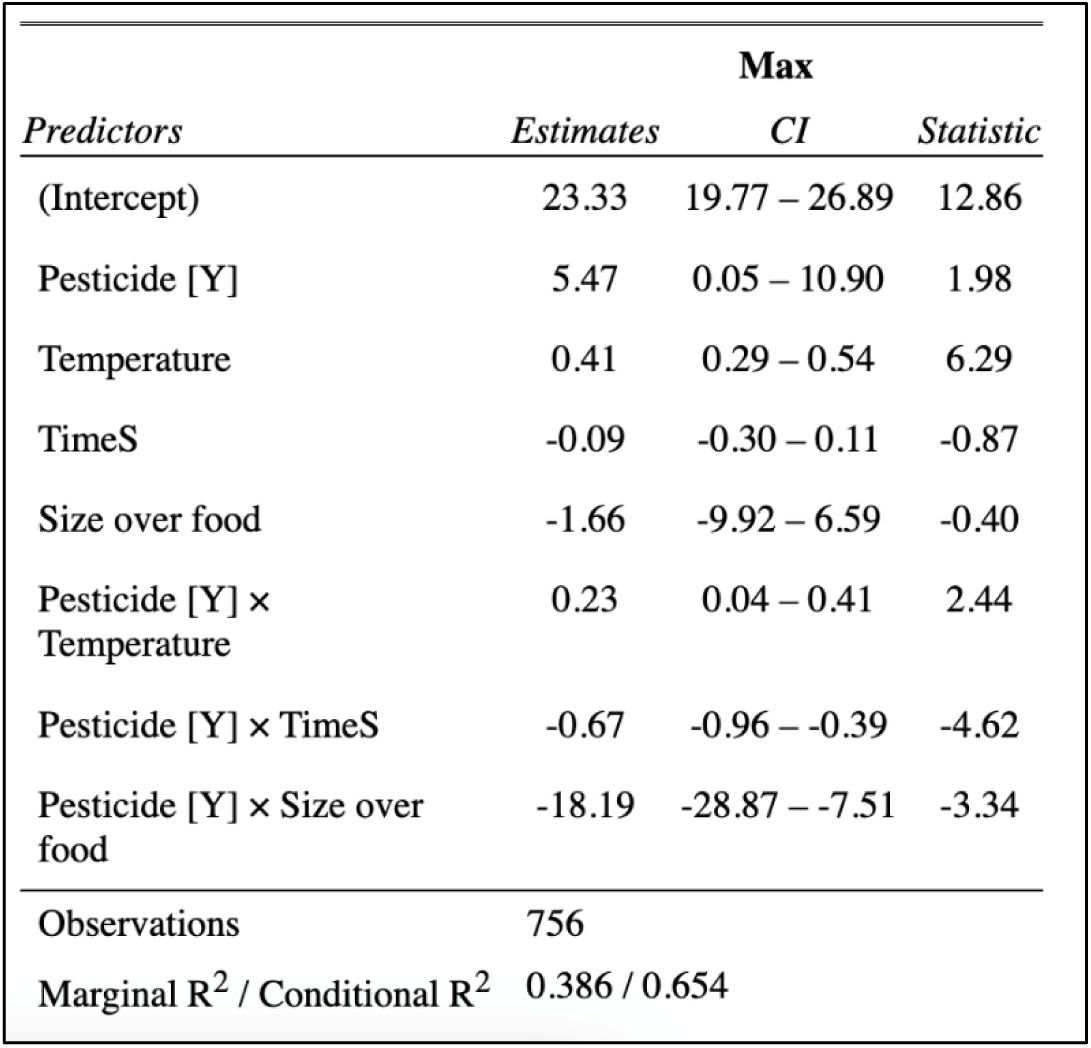
Output table from the statistical model predicting individual thorax temperature as a function of pesticide and temperature treatments whilst accounting for time and consumption per unit size (‘size over food’).

**Table S9:**
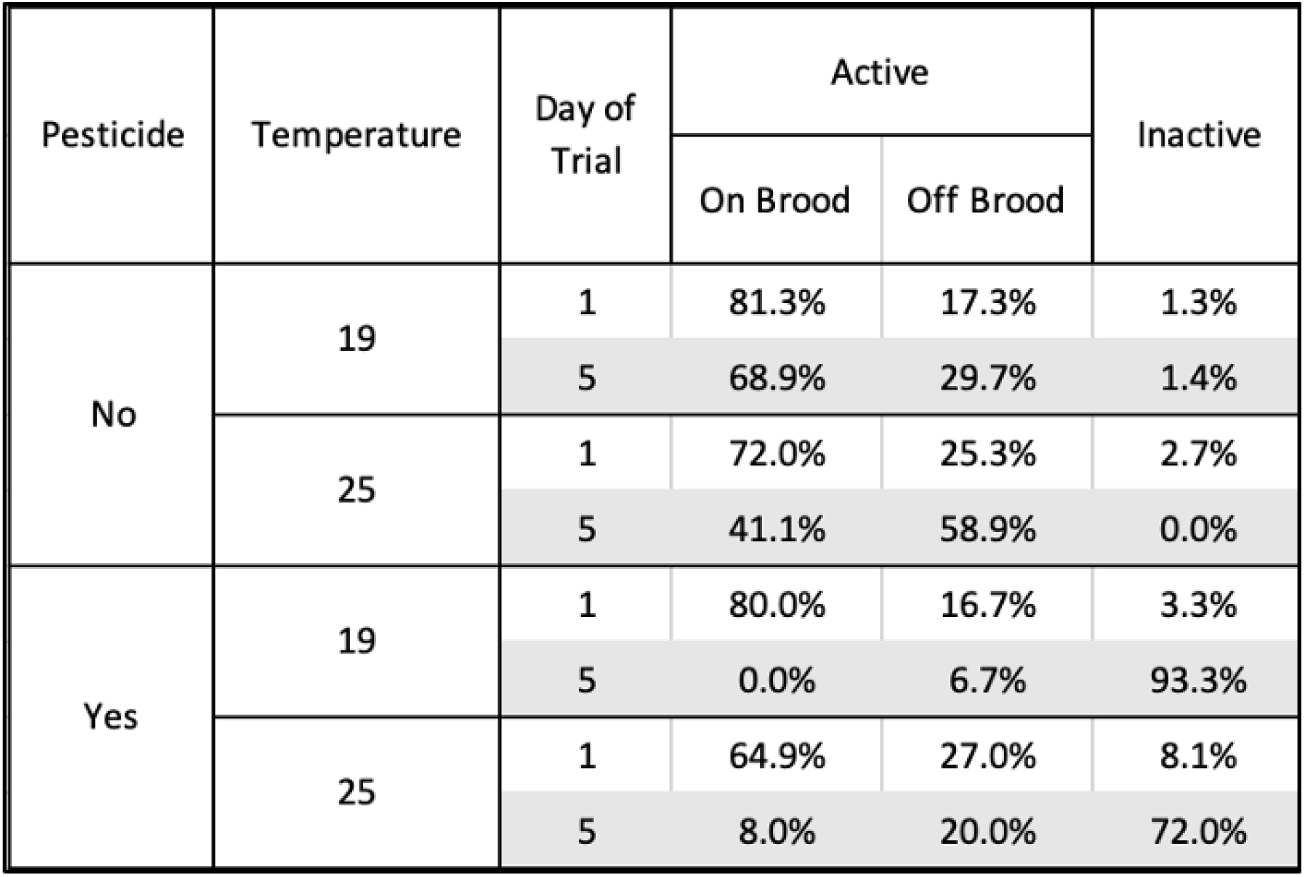
Percentage of bees that were active on the brood, active off the brood, and inactive when comparing between day 1 and 5 (final day) of the experiment (n = 300 snapshot assessments).

**Table S10:** Output table from the statistical model predicting the number of bees on brood as a function of pesticide and temperature treatments, accounting for time (‘time split’).

| *Predictors* | **Avg_on_brood** |  |  |
| --- | --- | --- | --- |
|  | *Estimates* | *CI* | *Statistic* |
| (Intercept) | 11.22 | 8.80 – 13.63 | 9.16 |
| Pesticide [Y] | 2.85 | -0.56 – 6.27 | 1.65 |
| Temperature [25] | 0.75 | -2.21 – 3.71 | 0.50 |
| time split | 0.62 | 0.40 – 0.84 | 5.48 |
| Pesticide [Y] ×<br>Temperature [25] | -2.28 | -6.46 – 1.90 | -1.08 |
| Pesticide [Y] × time<br>split | -1.05 | -1.36 – -0.75 | -6.72 |
| Observations | 185 |  |  |
| Marginal R<sup>2</sup> / Conditional R<sup>2</sup> | 0.345 / 0.561 |  |  |

**Table S11:** Output table from the statistical model predicting the proportion of bees on brood as a function of pesticide and temperature treatments and time (‘time split’).

| *Predictors* | **avg_proportion_bees_on_brood** |  |  |
| --- | --- | --- | --- |
|  | *Estimates* | *CI* | *Statistic* |
| (Intercept) | 1.00 | 0.70 – 1.31 | 6.52 |
| Pesticide [Y] | 0.50 | 0.25 – 0.76 | 3.87 |
| Temperature | -0.01 | -0.02 – 0.01 | -1.24 |
| time split | -0.01 | -0.06 – 0.03 | -0.53 |
| Pesticide [Y] ×<br>Temperature | -0.02 | -0.03 – -0.01 | -3.57 |
| Temperature × time split | 0.00 | -0.00 – 0.00 | 0.52 |
| Pesticide [Y] × time<br>split | -0.03 | -0.05 – -0.02 | -5.61 |
| Observations | 185 |  |  |
| R<sup>2</sup> / R<sup>2</sup> adjusted | 0.498 / 0.482 |  |  |

**Table S12:** Output table from the statistical model predicting brood temperature as a function of pesticide and temperature treatments and mediated by the average number of bees on the brood whilst accounting for time (‘time split mod’) and brood mass at the start of the trial (‘average pre brood mass’).

| *Predictors* | **Avg_hotspot** |  |  |
| --- | --- | --- | --- |
|  | *Estimates* | *CI* | *Statistic* |
| (Intercept) | 27.82 | 17.97 – 43.08 | 14.94 |
| Pesticide [Y] | 2.18 | 1.00 – 4.76 | 1.97 |
| Temperature | 1.01 | 0.99 – 1.03 | 1.05 |
| time split mod | 0.99 | 0.95 – 1.03 | -0.54 |
| Avg bees on brood | 1.00 | 0.98 – 1.03 | 0.21 |
| Pesticide [Y] ×<br>Temperature | 0.97 | 0.94 – 1.01 | -1.64 |
| Pesticide [Y] × time<br>split mod | 0.88 | 0.82 – 0.94 | -3.95 |
| Temperature × time split<br>mod | 1.00 | 1.00 – 1.00 | 0.35 |
| Pesticide [Y] × Avg bees<br>on brood | 0.95 | 0.91 – 0.99 | -2.36 |
| Temperature × Avg bees on<br>brood | 1.00 | 1.00 – 1.00 | -0.11 |
| (Pesticide [Y] ×<br>Temperature) × time split<br>mod | 1.00 | 1.00 – 1.01 | 2.64 |
| (Pesticide [Y] ×<br>Temperature) × Avg bees<br>on brood | 1.00 | 1.00 – 1.00 | 2.08 |
| Observations | 165 |  |  |
| R<sup>2</sup> Nagelkerke | 0.878 |  |  |

**Table S13:** Output table from the statistical model predicting brood temperature as a function of pesticide and temperature treatments and mediated by the average proportion of bees on the brood and whilst accounting for time (‘time split mod’) and brood mass at the start of the trial (‘average pre brood mass’).

| *Predictors* | **Avg_hotspot** |  |  |
| --- | --- | --- | --- |
|  | *Estimates* | *CI* | *Statistic* |
| (Intercept) | 28.38 | 16.95 – 47.51 | 12.70 |
| Pesticide [Y] | 0.73 | 0.35 – 1.53 | -0.83 |
| Temperature | 1.01 | 1.00 – 1.03 | 1.42 |
| time split mod | 0.99 | 0.92 – 1.06 | -0.24 |
| Avg proportion on brood | 1.00 | 0.65 – 1.53 | -0.01 |
| Pesticide [Y] × Temperature | 1.01 | 0.99 – 1.03 | 0.90 |
| Pesticide [Y] × time split mod | 0.95 | 0.86 – 1.04 | -1.08 |
| Temperature × time split mod | 1.00 | 1.00 – 1.00 | 0.25 |
| Pesticide [Y] × Avg proportion on brood | 1.24 | 0.72 – 2.11 | 0.78 |
| time split mod × Avg proportion on brood | 1.00 | 0.94 – 1.06 | -0.01 |
| (Pesticide [Y] × Temperature) × time split mod | 1.00 | 1.00 – 1.00 | 1.09 |
| (Pesticide [Y] × time split mod) × Avg proportion on brood | 0.96 | 0.89 – 1.03 | -1.21 |
| Observations | 165 |  |  |
| R<sup>2</sup> Nagelkerke | 0.874 |  |  |

**Table S14:** Eclosion of pupae and the number of adult bees counted during and at the end of the trial.

| Temperature | Pesticide | Visible Cocoons Change (Eclosion) (mean) | Total Bees During Experiment (mean) | Final Bee Counts (mean) |
| --- | --- | --- | --- | --- |
| 19 | N | 3.60 | 24.60 | 22.00 |
| 19 | Y | 0.20 | 17.80 | 14.40 |
| 25 | N | 5.75 | 26.25 | 26.25 |
| 25 | Y | 0.00 | 16.00 | 15.00 |

**Table S15:**
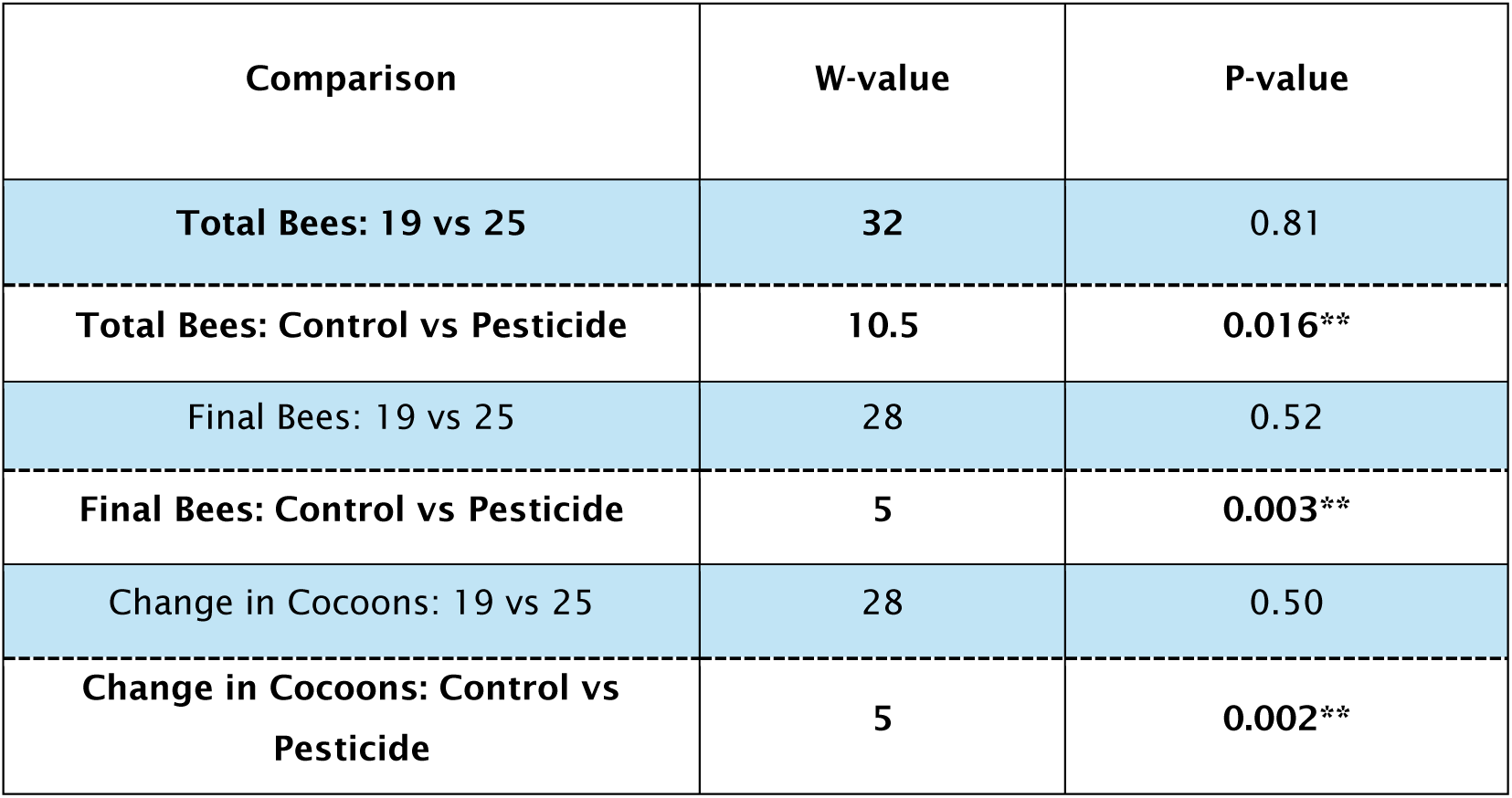
Wilcoxon test to compare the total adult bees between treatments.

**Table S16.** Output table from the statistical model predicting pupal development duration in experiments 3.

| Linear model predicting pupal development duration |  |  |  |
| --- | --- | --- | --- |
| *Predictors* | *Estimates* | *CI* | *Statistic* |
| (Intercept) | 13.3 | 12.462 – 14.052 | 33.2 |
| Temperature [Hot] | -5.24 | -6.252 – -4.234 | -10.4 |
| Pesticide [Y] | -0.791 | -1.933 – 0.352 | -1.38 |
| Start Age | -0.840 | -1.071 – -0.610 | -7.27 3 |
| Pesticide [Y] ×<br>Temperature | 0.928 | -0.340 – 2.196 | 1.46 1 |
| Num.Obs. | 77 |  |  |
| R2 | 0.817 |  |  |
| R2 Adj. | 0.807 |  |  |

**Table S17.** Sample size and treatments in the pupal rearing experiment n=106 at starting and n=77 that reached the final pupation stage.

| **Treatment** | **Individuals at start** | **Individuals that reached the final pupation stage** |
| --- | --- | --- |
| 25 °C Control | 21 | 12 |
| 25 °C Imidacloprid | 31 | 26 |
| 34 °C Control | 33 | 23 |
| 34 °C Imidacloprid | 21 | 16 |

**Table S18.** Model structure for experiment 3.

| Response Variable | Model Type | Link Function | Components | AIC | R<sup>2</sup> |
| --- | --- | --- | --- | --- | --- |
| Pupal development duration | lm | Identity | Temp * Exposure + StartAge | 268.42 | 0.817 |

